# Quantifying fluorescence lifetime responsiveness of environment sensitive probes for membrane fluidity measurements

**DOI:** 10.1101/2023.10.23.563572

**Authors:** Franziska Ragaller, Ellen Sjule, Yagmur Balim Urem, Jan Schlegel, Rojbin El, Dunja Urbancic, Iztok Urbancic, Hans Blom, Erdinc Sezgin

**Author notes:** Correspondence: Erdinc Sezgin 0046702318248.

## Abstract

The structural diversity of different lipid species within the membrane defines its biophysical properties such as membrane fluidity, phase transition, curvature, charge distribution and tension. Environment-sensitive probes, which change their spectral properties in response to their surrounding milieu, have greatly contributed to our understanding of such biophysical properties. To realize the full potential of these probes and to avoid misinterpretation of their spectral responses, a detailed investigation of their fluorescence characteristics in different environments is necessary. Here, we examined fluorescence lifetime of two newly developed membrane order probes, NR12S and NR12A, in response to alterations in their environments such as degree of lipid saturation, cholesterol content, double bond position and configuration and phospholipid headgroup. As comparison, we investigated lifetime sensitivity of the membrane tension probe Flipper in these environments. Applying fluorescence lifetime imaging microscopy (FLIM) in both model membranes and biological membranes, all probes distinguished membrane phases by lifetime, but exhibited different lifetime sensitivities to varying membrane biophysical properties (e.g. cholesterol). While the lifetime of Flipper is particularly sensitive to membrane cholesterol content, NR12S and NR12A lifetime is moderately sensitive to both cholesterol content and lipid acyl chains. Moreover, all probes exhibit longer lifetimes at longer emission wavelengths in membranes of any complexity. This emission-wavelength dependency results in varying lifetime resolution at different spectral regions, highly relevant for FLIM data acquisition. Our data provides valuable insights on how to perform FLIM with these probes and highlights both their potential and limitations.

## Introduction

The plasma membrane is a highly complex organelle comprising different lipid species, various membrane-associated proteins and glycocalyx components ^1^. This complexity ensures proper functionality of cellular mechanisms associated with the membrane such as cell signalling, intracellular membrane trafficking, endo- and exocytosis and cell division ^2^. The asymmetric distribution of the more than hundred different lipid species of diverse structure within the membrane bilayer is also essential for these cellular processes ^3,4^. Determined by high structural diversity, the collective membrane biophysical properties – membrane fluidity, phase transition, curvature, charge distribution and tension ^5,6^ – vary dynamically as a result of alterations in local membrane composition ^7^. Membrane biophysical properties and cellular processes are closely intertwined, emphasizing the need for further in-depth investigation of biophysical properties ^6,8^.

Investigating the influence of single structural changes (e.g., monounsaturated vs. saturated lipids) on biophysical properties in a cell plasma membrane poses a challenge due to the high complexity of the plasma membrane. To circumvent this problem, model membrane systems allowing for custom lipid composition are often exploited ^1,9,10^. To examine membrane biophysical properties such as polarity, hydration, viscosity and tension, among others, a variety of fluorescent probes have been developed in the past two decades, which respond to their associated environment by changes in intensity, emission wavelength or fluorescence lifetime ^11,12^. Solvatochromic membrane probes distinguish between membranes comprising saturated vs. unsaturated lipids by a shift of the emission maximum (towards longer wavelengths for membranes rich with unsaturated lipids) ^11^. The classical solvatochromic probe Laurdan ^13,14^, as well as other more recent probes such as di-4-ANEPPDHQ ^15^, NR12S ^16^, NR12A ^17^, NR4A ^17,18^ and Pro12A ^19^ have proven very useful for cell biology ^20^. Especially NR12S, NR12A and Pro12A are advantageous compared to previously synthesized probes due to increased brightness, larger emission shift upon changes in the environment, better defined plasma membrane location, decreased internalization and cytotoxicity as well as their suitability for live-cell imaging ^16,17,19,20^. Their spectral shift is examined using spectral or ratiometric imaging and subsequently quantified by calculation of the generalized polarization parameter (GP) or intensity ratios using spectral fitting ^21,22^ or fit-free spectral phasor approaches ^23^. Large red-shifts result in low GP-values and correlate with an increase in membrane fluidity, i.e. reporting on lower lipid order and decreased lipid packing ^14,21^. Of note, GP values or other ratios, do not directly disclose the underlying biophysical mechanisms causing the emission shifts. Further, a common misconception is that all solvatochromic probes sense the same membrane biophysical properties; as recent data shows that membrane probes Laurdan and di-4-ANEPPDHQ report on different membrane biophysical properties ^24^. Moreover, Pro12A, NR12S and NR12A exhibit varying sensitivities towards lipid saturation index, cholesterol content, configuration and position of lipid unsaturation and lipid headgroup, which is due to varying location and orientation of the probes in the membrane, resulting in subtle changes of their immediate environment ^25^.

Orthogonal to solvatochromic dyes, mechanosensitive planarizable push-pull probes (commercialized as Flipper-TR® and referred to as “Flipper” hereon) that incorporate into membranes have been developed ^26–28^. These probes report on their environment by changing their fluorescence lifetime ^26^. The lifetime is examined using fluorescence lifetime imaging microscopy (FLIM) and subsequently analysed by curve fitting, deconvolution or phasor lifetime methods ^29^. Mechanistically, Flipper probes are planarized by increasing physical compression (membrane tension), thereby generating stronger push-pull systems resulting in longer lifetimes ^26^. Flipper also senses increasing membrane order by an increase in lifetime and distinguishes liquid ordered (Lo) and liquid disordered (Ld) membrane phases ^27^. Other solvatochromic probes such as Laurdan ^30^, Nile Red ^31^ and di-4-ANEPPDHQ ^32^ among others also report on their environment by changes in their lifetime. Due to the intensity-independent nature, fluorescence lifetime measurements have advantages over intensity-based spectral measurements. Therefore, here, we aimed at investigating if the lifetime of NR12S and NR12A, as an intensity-independent readout, can be reliably used as a parameter to quantify changes in biophysical properties of the membrane. Following extensive characterization of NR12S and NR12A using spectral imaging ^25^, we thus quantitatively examined if there are differences in sensitivity between intensity-dependent emission shift and intensity-independent fluorescence lifetime. Furthermore, as comparison, the sensitivity of Flipper as lifetime probe to varying membrane biophysical properties was investigated. We determined the probes’ lifetimes in membranes of low complexity (large unilamellar vesicles (LUVs) and phase-separated giant unilamellar vesicles (GUVs)) and high complexity (cells and virus-like particles (VLPs)).

Overall, our results reveal that fluorescence lifetimes of NR12S and NR12A are mostly sensitive to high cholesterol content and distinguish membrane phases. The lifetime of Flipper is particularly sensitive to membrane cholesterol content, which may be related to an increase in membrane tension, and senses phase separation. Strikingly, all probes, especially NR12S and NR12A, exhibit longer lifetimes at longer emission wavelengths in membranes of any complexity. This emission-wavelength dependency of the fluorescence lifetimes is important to consider when performing FLIM experiments with these probes, i.e. selection of detection parameters. To conclude, our data provides valuable insights on how to perform FLIM with these probes, in model membranes as well as more complex systems, highlighting both their potentials and limitations to explore membrane biophysical properties.

## Material and Methods

### Materials

The following lipids and environment-sensitive probes were utilized: 1,2-diarachidonoyl-sn-glycero-3-phosphocholine (DAPC, 20:4/20:4 PC), 1,2-dipetroselenoyl-sn-glycero-3-phosphocholine (Δ6cis DOPC, 18:1/18:1 PC), 1,2-dioleoyl-sn-glycero-3-phosphocholine (Δ9cis DOPC, 18:1/18:1 PC), 1,2-dielaidoyl-sn-glycero-3-phosphocholine (Δ9trans DOPC, 18:1/18:1 PC), 1-palmitoyl-2-oleoyl-glycero-3-phosphocholine (POPC, 16:0-18:1 PC), 1,2-dipalmitoyl-sn-glycero-3-phosphocholine (DPPC, 16:0/16:0 PC), 1-palmitoyl-2-oleoyl-sn-glycero-3-phospho-L-serine (POPS, 16:0/18:1 PS), 1-palmitoyl-2-oleoyl-sn-glycero-3-phosphoethanolamine (POPE, 16:0/18:1 PE), cholesterol, brain octadecanoyl sphingomyelin (SM, 18:0) (Avanti Polar Lipids), Flipper-TR® ^27^, NR12S ^16^ and NR12A ^17^. NaCl and HEPES were obtained from Sigma-Aldrich (St. Louis, MO, USA). PBS, high-glucose DMEM and Leibovitz’s L15 medium were acquired from ThermoFisher Scientific.

### Large unilamellar vesicle preparation and staining

For LUV production, the desired lipid mixture was prepared in chloroform (0.5 mg/ml). After removal of the solvent by nitrogen flow, the lipid film was hydrated with 1 ml buffer (150 mM NaCl, 10 mM HEPES, pH 7.4) and the solution was vortexed vigorously to disperse the lipid into the buffer. The solution was sonicated using a tip sonicator (power 3, duty cycle 40%) for 10 min. LUVs were stored under nitrogen at 4 °C. The LUVs were stained with 1 μM of Flipper, NR12S or NR12A. As control (no environment-sensitive dye) 1μM AlexaFluor 488 in water was used. All bulk samples were imaged in μ-Slides (18 well glass bottom, ibidi), previously blocked with 3 mg/ml BSA in PBS, at room temperature.

### Giant unilamellar vesicle preparation and staining

Phase-separated GUVs (SM:DOPC:Chol 2:2:1) were prepared according to a previously described protocol ^9^. Using custom-built GUV Teflon chambers with two platinum electrodes, GUVs were generated by electroformation ^33^. A volume of 6 μl of lipid dissolved in chloroform (1 mg/ml total lipid concentration) was homogeneously distributed on the electrodes, dried under nitrogen stream and placed in 300 nM sucrose solution (370 μl). Electroformation was performed at 2 V and 10 Hz at 70 °C (above the specific lipid transition temperature) for 1 h, followed by 2 V and 2 Hz for 30 min. GUVs were stained at a final concentration of 300 nM Flipper and 100 nM of NR12S or NR12A. The GUVs were imaged in μ-Slides (18 well glass bottom, ibidi), previously blocked with 3 mg/ml BSA in PBS, at room temperature.

### Cell maintenance and staining

NRK 52E, U2OS, RBL and HEK293T cells were cultured in DMEM (high glucose, without pyruvate) with 10 % FBS at 37 °C and 5 % CO_2_. HEK293T cells were utilized for VLP preparation. For imaging, 1×10^4^ cells (NRK 52E, U2OS and RBL) per well were seeded in μ-slides (18 well glass bottom, ibidi). Cells were stained with 1 μM of Flipper, NR12S and NR12A in phenolred- and serum-free L15 medium. The cells were not washed before imaging.

### Preparation of virus-like particles and staining

VLPs were produced as described previously ^34^ with small modifications. To produce pseudotyped non-fluorescent VLPs, HEK293T cells were seeded at a confluency of ∼70 % in T75 flasks and co-transfected 6 hours later with 7.5 µg of the lentiviral packaging vector psPAX2 (gift from Didier Trono - Addgene plasmid # 12260) and 15 µg of the plasmid encoding the respective viral surface protein (pCMV14-3X-Flag-SARS-CoV-2 S was a gift from Zhaohui Qian - Addgene plasmid # 145780; delta spike expression plasmid kindly provided by Benjamin Murrell; Ebola GP expression plasmid kindly provided by Jochen Bodem) using Lipofectamine™ 3000 (ThermoFisher) according to the manufacturer’s recommendations. After 12 h, medium was replaced by phenolred-free DMEM supplemented with 10 % FCS and VLPs harvested twice after 24 h. VLP-containing supernatant was sterile filtered through a 0.45 µm PES filter and fiftyfold enriched using LentiX concentrator following the manufacturer’s protocol (Takara). The VLPs were diluted 1:1 with PBS and stained at a final concentration of 300 nM Flipper or 100 nM NR12S or NR12A before imaging in μ-Slides (18 well glass bottom, ibidi), previously blocked with 3 mg/ml BSA in PBS at room temperature. Of note, possible contamination by other particles of similar size and density such as lipoproteins or extracellular vesicles (EVs) cannot be excluded with the VLP preparation protocol utilized, possibly influencing the lifetime measurements by masking potential differences between VLP species.

### Fluorescence lifetime imaging microscopy

All FLIM measurements were performed on a Leica SP8 3X STED with FALCON FLIM/FCS, utilizing a STED white HC PL APO CS2 100x/1.40 oil objective. Excitation output power was set to 70 % of a pulsed white-light-laser and software-tuned to optimal settings for each experiment. All probes were excited at 488 nm and emission was collected within the 500-700 nm range through prism-based spectral selections either in intervals of 20 nm (sequentially) for LUVs and VLPs, or within 500-600 nm and 600-700 nm for phase-separated GUVs and cells. Emission was collected using a SMD HyD detector set to photon counting mode (10% internal gain). Power settings for FLIM ensured a maximum count rate of 0.5 photons per laser pulse and all measurements were taken at 20 MHz repetition rate, except when investigating laser frequency influences where measurements at 40 and 80 MHz were performed. For more detailed information on specific acquisition parameters see Supplement Table 1.

### Fluorescence lifetime analysis

Lifetime analysis was performed using the Leica Application Suite LAS X FLIM/FCS software (version 4.5.0). All fluorescence decay curves were analysed by n-exponential reconvolution fitting using the instrument response function (IRF) calculated by the FALCON FLIM software within the range 0.2-45 ns (20 MHz), 0.2-25 ns (40 MHz) and 0.2-12.5 ns (80 MHz). For all analyses we further used an intrinsic standard (high speed) photon filter, ensuring that only single photons detected between two laser pulses are used for the lifetime decay ^35^. For LUVs, cells and VLPs whole image analysis was performed (histograms of photons pooled from the whole image were fitted), whereas for phase-separated GUVs histograms of photons from manually selected field-of-views were fitted (Supplement Figure 4). Different selected emission windows (i.e. width of 20 or 100 nm) were fitted individually, except for comparison of the decay rates of the probes in Δ9cis DOPC vs. DPPC:Chol 50:50 LUVs (Figure 1B), where the decays across detection windows were combined (200 nm emission window). For the comparison of intensity-based vs. lifetime-based analysis (Figure 5), the decays across detection windows within the range of 500-600 nm or 500-700 nm were combined, to ensure best lifetime resolution (according to Figure 1E and Supplement Figure 1). Fluorescence decays of less than 10^4^ photons were excluded due to rendering unreliable fitting analysis ^29,36^. Technical replicates exhibiting unrealistic lifetime values or inappropriate fits were identified as outliers and excluded from the plots. For more detailed information of specific fitting parameters see Supplement Table 2, especially for the selections of components used for n-exponential reconvolution fitting. All lifetime values shown in the plots correspond to the ‘mean intensity weighted lifetime’ calculated by the LAS X FLIM/FCS software Using Equation 1:

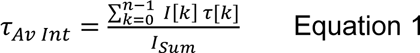

**Figure 1:**
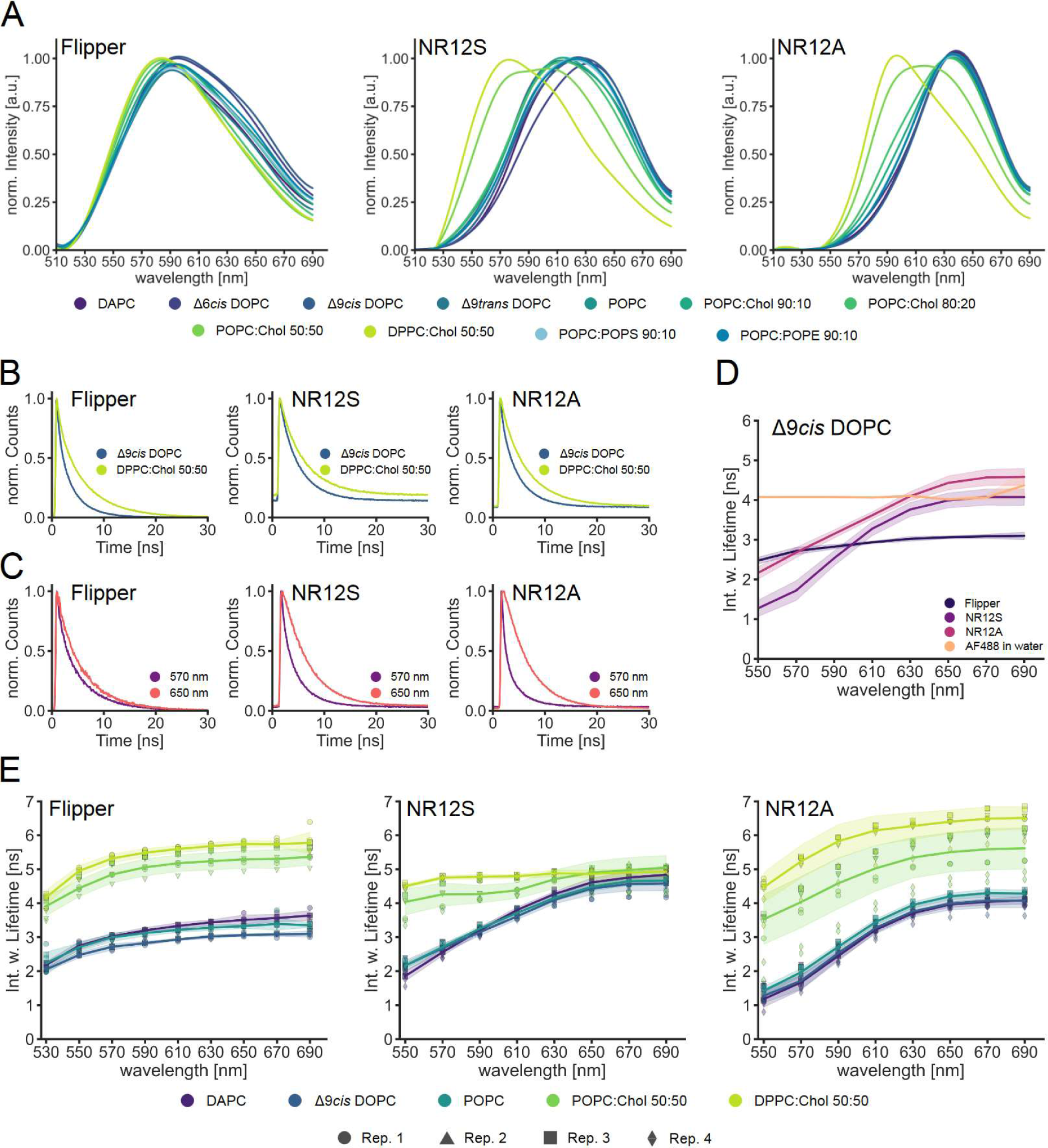
Characterization of Flipper, NR12S and NR12A in different lipid environments. Spectral fluorescence lifetime measurements of the probes in LUVs were carried out from 500-700 nm in intervals of 20 nm. Multiexponential curve fitting was performed for the fluorescence decays (for details see Material and Methods). A| Normalized intensity spectra of Flipper (left), NR12S (middle) and NR12A (right) in varying lipid environments. B| Normalized fluorescence decays (full spectrum, 500-700 nm) of Flipper (left), NR12S (middle) and NR12A (right) in Δ9*cis* DOPC or DPPC: Chol 50:50. C| Normalized fluorescence decays at 570 nm vs. 650 nm of Flipper (left), NR12S (middle) and NR12A (right) in POPC:Chol 80:20. D| Spectrally resolved intensity weighted lifetime of Flipper, NR12S and NR12A in Δ9*cis* DOPC and the control AF488 in water. Line corresponds to the median of individual biological replicates (n=3 or 4). Band corresponds to standard deviation. E| Spectrally resolved intensity weighted lifetime of Flipper (left), NR12S (middle) and NR12A (right) in different lipid environments with varying saturation index. Line corresponds to the median of individual biological replicates shown with different symbols (n=3 or 4). Band corresponds to standard deviation.

With τ*_Av Int_* being the ‘mean intensity weighted lifetime’, *I* being the intensities associated with each exponential component, normalized to the time resolution of the measured decay curve, τ being the lifetime, and *I_Sum_* being the sum of fluorescence intensity for all components.

### Calculation of normalized intensity spectra and GP

After determination of the lifetime values all further analysis was carried out using Python 3.10. To calculate normalized intensity spectra, the sum intensity (*I_Sum_*) obtained from the FLIM measurements were normalized to their maximum values. This was performed for each 20 nm emission window recorded of all probes across different lipid compositions. For better visualization, the normalized intensity values were interpolated using a cubic spline function.

To compare the resolution of lifetime and intensity of the probes in different lipid environments the generalized polarization (GP) parameter was calculated using Equation 2:

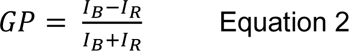

with *I_B_* and *I_R_* being fluorescence signal intensities at blue-or red-shifted emission wavelengths, respectively, for liquid ordered, λ_Lo_, and liquid disordered phases, λ_Ld_. The GP can adopt values ranging between +1 and −1, according to Equation 1. The GP value is a ratiometric (relative) quantification depending on selected λ_Lo_ and λ_Ld_ values as well as on the environment-sensitive probe. The respective sum intensity counts obtained from the FLIM measurements were used to calculate the median GP value (three replicates) of the investigated probes in different lipid environments. The median GP values were then compared with the median mean intensity weighted lifetime values (same three replicates) obtained within 500-600 nm or 500-700 nm. For spectral imaging analysis (∼9 nm channels), we previously concluded selection of λ_Lo_ and λ_Ld_, 557 and 664 nm for NR12S, and 583 and 673 nm for NR12A, respectively ^25^. At these wavelengths, the differences in intensity of ordered and disordered phases generated the largest GP range, while maintaining sufficient fluorescence signal. Correspondingly, for the spectrally resolved FLIM data, we selected similar detection wavelengths (λ_Lo_ and λ_Ld_) for GP calculations, i.e. 550 and 670 nm for NR12S, and 590 and 670 nm for NR12A. For Flipper 570 and 650 nm were chosen.

### Phasor analysis

Phasor analysis of lifetime was performed using the LAS X FLIM/FCS software (version 4.5.0). To display the detected and time sorted photons in the phasor plot a wavelet filter with threshold 5 was applied, for better differentiation of the phasor photon clouds. For analysis, the centre of the phasor-photon clouds was manually selected with the circular selection tool of a radius 20 (see Figure 2B). The average estimated fluorescence lifetime within the circle was used for further comparisons.

**Figure 2:**
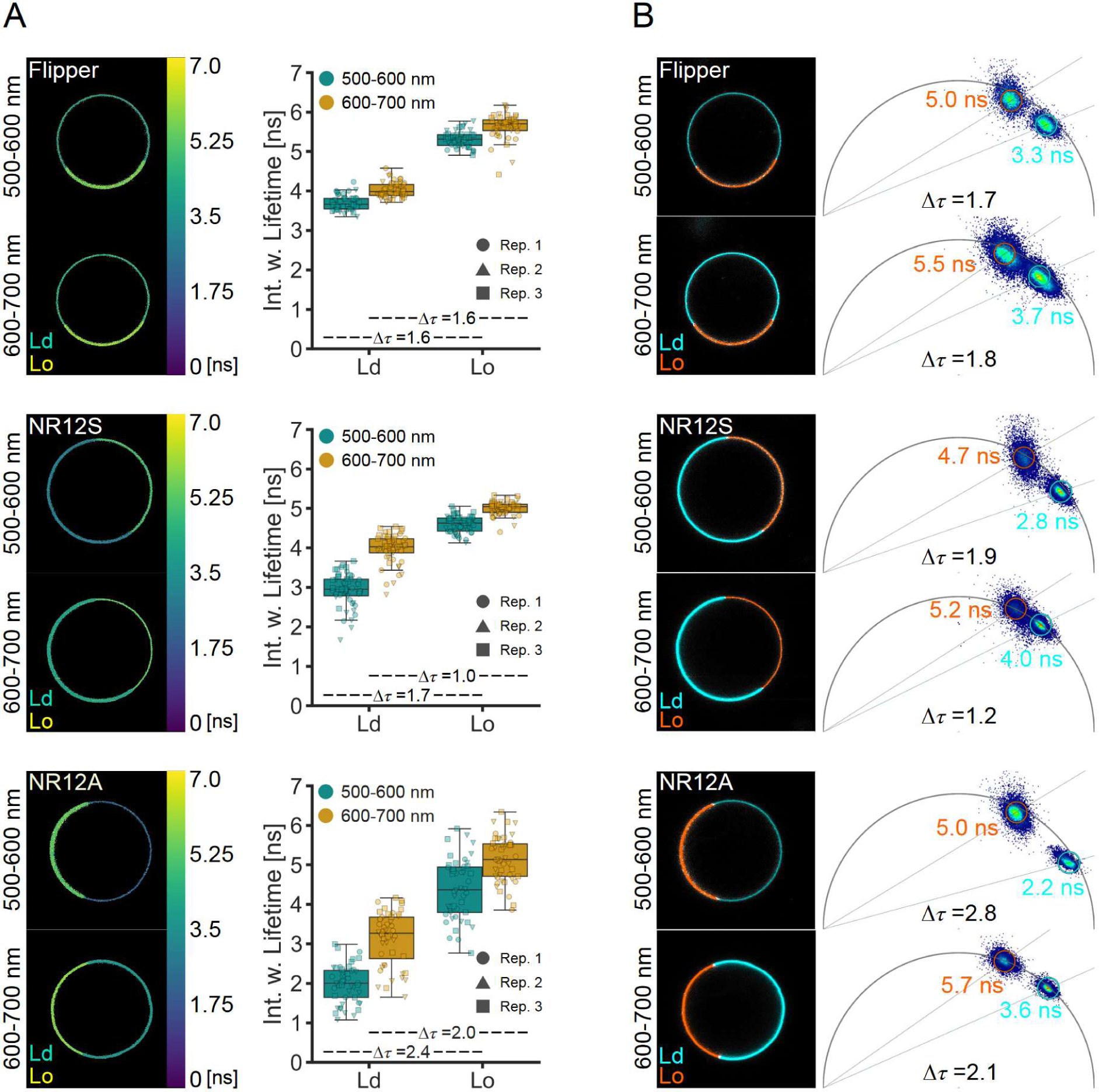
Lifetimes of Flipper, NR12S and NR12A are sensitive to phase-separation. Lifetime measurements of the probes in phase-separated GUVs were carried out at 500-600 nm or 600-700 nm emission. Multiexponential curve fitting was performed for the fluorescence decays (for details see Material and Methods). A| Example images of lifetime-colour coded phase-separated GUVs and comparison of the intensity weighted lifetime in liquid-disordered (Ld) vs. liquid-ordered (Lo) phase at different emission wavelengths of Flipper (above), NR12S (middle), NR12A (below). Different symbols correspond to individual biological replicates (n=3). Δτ values were calculated from the mean of the intensity weighted lifetimes. B| Phasor analysis of Flipper (above), NR12S (middle), NR12A (below). Example images of lifetime separation (left) according to the clouds on the phasor plot (right) with Ld shown in cyan and Lo in orange. Δτ values were calculated from the average lifetimes of the phasor clouds.

## Results and Discussion

Spectral imaging has shown that environment sensitive fluorescent probes NR12S and NR12A have individual sensitivities to membrane biophysical properties such as saturation index, cholesterol content, double bond position and configuration as well as lipid headgroup charge and geometry ^25^. The shift in the probes’ emission spectrum in different lipid environments reflects these sensitivities (Figure 1A). Although Flipper is a lifetime sensitive probe, it also exhibits minor shifts towards longer wavelengths of its emission spectrum in more disordered lipid environments (Figure 1A), which is in line with previous studies ^26^. As NR12S and NR12A are suitable for investigation of lipid environments by spectral imaging, we aimed to examine their suitability for fluorescence lifetime imaging analysis. Therefore, we characterized their lifetime behaviour alongside Flipper probe with FLIM.

### Lifetime increases at longer emission wavelengths

Regarding the fluorescence decays of all the probes in a disordered vs. ordered (Δ9*cis* DOPC vs. DPPC:Chol 50:50 LUVs) lipid environment, we observed a shift towards longer decay times in the ordered lipid environment (Figure 1B). Interestingly, we also observed a shift of the fluorescence decay towards longer times for all three probes in LUVs of the same lipid composition at longer wavelengths (Figure 1C). This shift is more pronounced in NR12S and NR12A compared to Flipper. Due to this observation, we further investigated the fluorescence decay time in 20 nanometre wavelength intervals from 500-700 nm (laser repetition rate 20 MHz), to quantitatively evaluate the changes in lifetime at different emission wavelengths. We obtained the intensity weighted fluorescence lifetime of all probes (from here on referred to as lifetime) by exponential reconvolution fitting of the decay times (for details on data acquisition and curve fitting analysis, see Methods and Supplement Table 1). The lifetimes of all three membrane probes in Δ9*cis* DOPC increase with longer emission wavelength, while a common control dye (AlexaFluor 488 in water) does not exhibit any such difference as a function of collection window (Figure 1D). The increase in lifetime is less pronounced for Flipper, reaching a plateau already at around 600 nm compared to the lifetimes of NR12S and NR12A, which plateau at 650 nm. The observed emission-wavelength dependency of the lifetimes of all probes is most likely caused by solvent relaxation. Upon excitation, the dipole moment of the fluorophore is rapidly reoriented resulting in an energetically unfavourable Franck-Codon state ^37^. The molecules of the solvation envelope of the fluorophore then reorient, allowing the system to relax and thereby lowering the energy of the excited state. This reorientation of the solvent envelope takes time, which is reflected in progressively increasing the emission red-shift. The longer the fluorophore remains in the excited state, the longer the wavelength of the photons it emits ^37^. By splitting the overall fluorescence decay of the probes in 20 nm intervals in LUVs, we collected photons of short wavelength with short lifetimes from not fully relaxed fluorophores, which contrast photons of long wavelength with long lifetimes, from fully relaxed fluorophores. The emission of Flipper is less affected by solvent relaxation ^26^ compared to NR12S and NR12A as confirmed by detecting only minor shifts in its emission spectra in different lipid environments (Figure 1A). The detected emission wavelength dependency of the lifetimes was previously reported for Nile Red in different solvents ^31^.

### Flipper lifetime is sensitive to cholesterol content but not acyl chain order

We then examined the lifetimes of Flipper, NR12S and NR12A in different lipid environments of varying cholesterol content, saturation index, double bond position and configuration, as well as headgroup to investigate and quantify the probes’ lifetime variability and sensitivity. The emission wavelength dependency of the lifetime of Flipper was observed in all lipid environments (Figure 1E and Supplement Figure 1). Flipper was shown to be able to differentiate between mono-unsaturated and saturated lipids (POPC:Chol 50:50 and DPPC:Chol 50:50) by an increase in lifetime (Figure 1E). However, among poly-or mono-unsaturated lipids (DAPC, Δ9*cis* DOPC and POPC) the lifetime changes for Flipper are much smaller. Double bond position and configuration (Δ6*cis* DOPC, Δ9*cis* DOPC and Δ9*trans* DOPC) as well as headgroup charge and geometry (POPS and POPE) could hardly be differentiated by lifetime analysis (Supplement Figure 1). However, increasing amounts of cholesterol in the membrane result in increasing lifetime, which is in line with a previous study ^27^. Lipid packing and membrane tension are closely connected, as changed lipid composition (e.g. cholesterol introduction) as well as applied tension (e.g. micropipette aspiration) strongly influence lipid packing ^5,27^. This renders it almost impossible to distinguish the effect of membrane tension or lipid composition on the change of the lifetime of Flipper in model membrane systems ^27^, showing that fluorescent probes can be sensitive to multiple biophysical parameters.

### Lifetimes of NR12S and NR12A are highly dependent on emission wavelength

Next, the lifetime sensitivity of NR12S and NR12A in different lipid environments was examined. The lifetime of NR12S increases in presence of high cholesterol content and upon switching from monounsaturated to saturated lipids (POPC:Chol 50:50 and DPPC:Chol 50:50) (Figure 1E and Supplement Figure 1). The sensitivity to cholesterol is in line with previous results both in model membranes (LUVs) ^38^ and more complex membranes (neuronal membranes of the hippocampus) ^39^. Interestingly, these two lipid compositions are the only ones in which emission-wavelength dependency of the lifetime of NR12S was much less pronounced. While the lifetime differences of NR12S at shorter emission wavelengths (550-610 nm) are quite prominent, they are much smaller at longer emission wavelengths (630-690 nm). Similar to NR12S, NR12A can only distinguish lipid compositions of high cholesterol content and saturated from monounsaturated lipids from the other lipid compositions (Figure 1E and Supplement Figure 1). Among the three probes, the lifetime of NR12A shows the most pronounced emission-wavelength dependency: while the lifetime of NR12A can distinguish the two high cholesterol lipid compositions across the entire emission spectrum, the sensitivity is better (larger lifetime changes) at shorter emission wavelength regions (550-610 nm). As mentioned above, the strong emission-wavelength dependency of NR12S and NR12A are due to solvent relaxation. Additionally, Nile Red-derived dyes exhibit complex photophysical behaviour including a twisted internal charge transfer (TICT) ^31,40,41^, which competes with solvent relaxation causing multiple emitting species affected by the TICT with varying fluorescence properties, as observed in time resolved emission spectrum (TRES) experiments of NR12S and NR12A ^25^. Of note, fitting of the decay curves of NR12S and NR12A at longer wavelength regions (630-690 nm) was less accurate compared to Flipper indicated by the χ^2^ values (Supplement Figure 2).

### Optimal laser repetition rate is crucial for FLIM measurements

To investigate the effect of parameter fitting we also studied the hardware performance in our FLIM system. Notably, the frequency of the pulsed laser has a big impact on reliable lifetime values. We observed that laser excitation frequencies of 40 or 80 MHz cause an overestimation of fluorescence lifetimes and less reliable fitting (indicated by χ^2^ values) especially in the longer emission wavelength region and more ordered lipid compositions (NR12A as example, Supplement Figure 3A). Looking more closely at the fluorescence decay curves and the fitting performance, the offset increases with rising frequency as the decay is not completed within one laser pulse, and the accuracy of the fit decreases indicated by the increasing residuals around the laser pulse (NR12S in DPPC:Chol 50:50 as example, Supplement Figure 3B).

In lifetime analysis (via fitting time correlated single photon counting histograms), a pile-up effect occurs at high photon count rates, where photons with short arrival times are overrepresented due to the detector dead-times, resulting in an overall underestimation of lifetimes ^29^. We therefore strongly recommend selecting a low laser repetition rate (e.g., 20 MHz) that allows for complete fluorescence decays and for intervals between pulses that are around 10 times longer than the average lifetime, including fluorescence emission detection rates of less than one photon per pulse. Our conclusions are also in line with the study examining the laser frequency influence on the lifetime of Flipper in more detail ^42^.

### Lifetimes of NR12A and NR12S distinguish membrane phases better at the green spectral region

Lateral heterogeneity within membranes is crucial for their varying functions ^6^. While disordered phases are characterized by mostly unsaturated lipids, ordered phases are enriched in saturated lipids and cholesterol ^5^. As revealed above, all probes show lifetime sensitivity to high cholesterol content, and we thus wanted to examine whether the emission-wavelength dependency of the lifetime persists in other membrane systems and whether liquid-ordered (Lo) from liquid-disordered (Ld) phases are separated equally well at different spectral regions.

Therefore, we measured lifetimes of the probes in phase-separated GUVs (sphingomyelin:DOPC:Chol 2:2:1). Although spectrally fine-resolved lifetime measurements give a more detailed insight on plasma membrane properties, the approach was not feasible for the use in larger model membrane systems or cells due to collection constraints of detected photons (i.e. photon budget). For reliable multi-component exponential fit analyses, a high number of detected photons is crucial (10^2^, 10^4^ and 10^6^ for mono-, bi- and tri-exponential fitting) ^29,36^ in each emission window, which requires high laser power, multiple frame repetitions and almost completely immobile samples. Instead of narrow spectral bands, we therefore collected photons in two wider channels of equal wavelength, separating the extreme peaks in the emission spectrum (Figure 1A): at shorter (550±50 nm; referred to as green) and at longer (650±50 nm; referred to as red) wavelengths (Figure 2A). Moreover, to minimize the contribution of noise and obtain the needed photon counts for fitting analysis, we selected regions of interest (ROIs) of the Lo and Ld phase separately, instead of performing pixel-wise fitting (Supplement Figure 4A). The same selection was used for lifetime analysis in the green and red channel. All three probes can differentiate Lo from Ld phases by an increase in lifetime in both emission windows, with higher lifetimes in the red (Figure 2A), confirming the experiments in LUVs discussed above. Flipper differentiates the phases equally well in both emission windows with a shift in lifetime of Δτ=1.6 ns, which is in line with previous investigations ^27^ and in silico studies ^43^. The Lo labelling of Flipper seems to be more prominent, which has been previously explained by the increased oscillator strength in its more planarized form, which yields more photons ^26^. In contrast, NR12S and NR12A exhibit a higher lifetime resolution for the membrane phases in the green with Δτ=1.7 ns and Δτ=2.4 ns compared to Δτ=1.0 ns and Δτ=2.0 ns in the red, respectively. This is in line with our experiments performed in LUVs. At shorter wavelengths, NR12S and NR12A showed higher Δτ between Lo and Ld, however the heterogeneity of their lifetime is higher than of Flipper. Fitting of the decay curves according to the χ^2^ values was quite reliable for all three probes with some outliers occurring for NR12S and NR12A (Supplement Figure 4B).

### Phasor analysis of lifetime distinguishes membrane phases better at the green spectral region

As a fit-free technique, phasor-analysis of fourier-transformed fluorescence decays generates lifetime maps of pixel detected photon arrival times in a 2D phasor plot ^29,44^. Fluorescent probes with a mono-exponential decay locate on the universal semicircle in these plots. Whereas probes with multi-exponential decaying properties locate within the universal semicircle ^44^.

To complement our decay fitting approach, we employed phasor analysis to investigate the lifetime of the probes in Lo and Ld phases (Figure 2B). First, the location of the phasor clouds indicate the lifetime complexity: Flipper locates within the semi-circle irrespectively of membrane phase or detection window, indicating a multi-exponential decay in line with the performed biexponential curve fitting and previous studies ^27^. NR12S locates within the semi-circle at shorter wavelengths, but locates on the edge of the semi-circle at longer wavelengths, indicating a multi-exponential decay at shorter and a mono-exponential decay at longer wavelengths. For NR12A only the Lo phase at longer wavelengths is located on the semi-circle indicating a mono-exponential decay. All other phasor clouds are located within the semi-circle, thus indicating multi-exponential decays. The observation of the distribution of mono- and multiexponential decays of NR12S and NR12A agrees with the exponential curve fitting analysis (Supplement Table 2). Previous studies with NR12S used a biexponential fitting analysis, in line with our results ^38,39^. Concerning fit-free phasor analysis, all probes can easily separate the phases as seen by two distinct clouds in the phasor plots. The shift in lifetimes obtained from the phasor analysis follow a similar trend in both spectral windows as seen above, further confirming the emission-wavelength dependency. NR12S and NR12A again show better separation at the green spectral region, as shown by Δτ values (inset phasor plots Figure 2B).

### Lifetime imaging in cells confirms wavelength dependency

Next, to evaluate the applicability of these probes with lifetime analysis in complex cellular environments, we used different adherent cell lines NRK 52E, U2OS and RBL (Figure 3). Again, due to photon budget constraints, photons were collected in the broader green and red channels, similar to the phase-separated GUVs. Given that the membrane probes are fluorogenic (i.e. only fluoresce upon integration into the membrane ^11^), the cells were imaged directly after staining to avoid internalization during image acquisition. Longer incubation times with Flipper have shown internalization into endocytic compartments, displaying a shorter lifetime, thus affecting the overall lifetime in whole image analysis ^42^, which is why we kept incubation times short (see Methods). The emission-wavelength dependency with longer lifetimes at longer wavelengths was again observed for all three probes in line with the results in model membranes. Flipper and NR12S do not show large lifetime shifts among the different cell types (Figure 3A, B).

**Figure 3:**
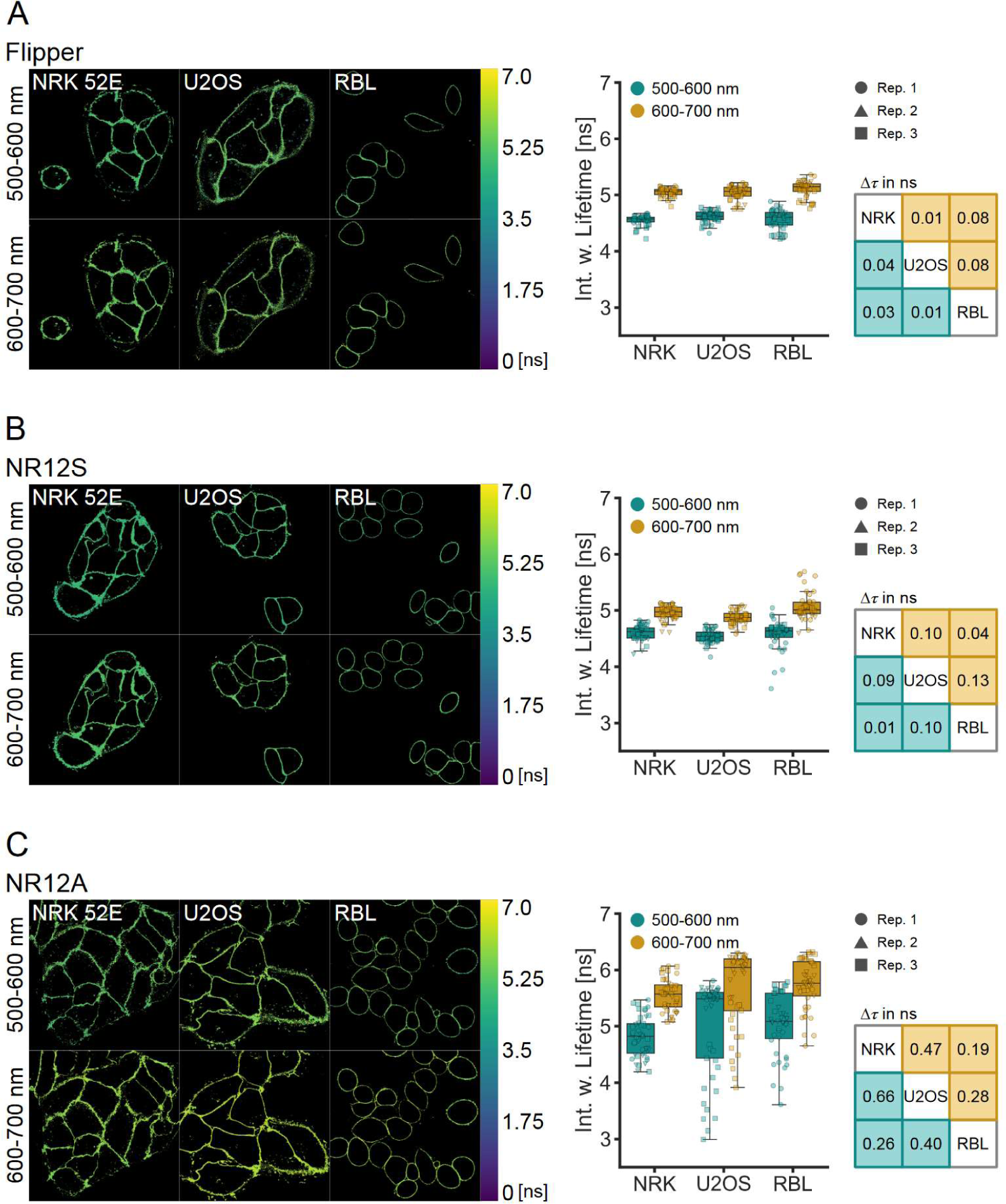
Lifetimes of Flipper, NR12S and NR12A in different cell types. Lifetime measurements of the probes in NRK 52E, U2OS and RBL cells were carried out at 500-600 nm and 600-700 nm emission windows. Multiexponential curve fitting was performed for the whole-image fluorescence decays (for details see Material and Methods). A| Example images of lifetime-colour coded cells stained with Flipper and comparison of the intensity weighted lifetime in different cell types and at different emission wavelengths. Different symbols correspond to images of individual biological replicates (n=3). Table indicates Δτ values of the medians between the cell types in the two channels. B| Example images of lifetime-colour coded cells stained with NR12S and comparison of the intensity weighted lifetime in different cell types and at different emission wavelengths. Different symbols correspond to images of individual biological replicates (n=3). Table indicates Δτ values of the medians between the cell types in the two channels. C| Example images of lifetime-colour coded cells stained with NR12A and comparison of the intensity weighted lifetime in different cell types and at different emission wavelengths. Different symbols correspond to images of individual biological replicates (n=3). Table indicates Δτ values of the medians between the cell types in the two channels.

Similar to lifetime measurements in LUVs and GUVs, Flipper lifetime shifts (Δτ values) are within the same range at shorter and longer emission wavelengths. This is also the case for NR12S, contrasting experiments in simpler lipid environments, where the lifetime resolution was better at shorter wavelengths. Lifetime analysis of NR12A resulted in very heterogenous lifetime values for U2OS and RBL cells, which obscure resolving differences in membrane biophysical properties (Figure 3C). Fitting analysis for NR12A was less reliable compared to Flipper and NR12S indicated by increasing χ^2^ values, especially at longer detected emission wavelengths (Supplement Figure 5), which could explain the lifetime heterogeneity. To obtain sufficient photon counts for multi-exponential curve fitting of the decay and avoid phototoxicity due to long acquisition times in live cells, we performed whole image analysis instead of pixel-wise fitting. This approach may on the other hand average out small existing differences, emphasizing a limitation of FLIM analysis with these probes in complex environments.

### Lifetimes of Flipper and NR12A distinguish delta-Spike VLPs

Virus envelope is a complex structure where environment-sensitive probes are helpful to understand biophysics of host-pathogen interactions. Virus-like particles have wide application in biology and medicine, ranging from unravelling viral structures, investigation of virus-host cell interactions to vaccine development ^45^. Viruses with a lipid envelope obtain the lipids from the host cell plasma membrane upon virus budding. During the budding process lipid-protein interactions drive the accumulation of certain lipids into the viral envelope ^46^ resulting in varying envelope lipid composition of different viruses with potentially distinctive biophysical properties. Especially, SARS-CoV-2 Spike variants evolved to have increased positive charge resulting in enhanced binding ^47^. Whether these severe surface changes are accompanied by alterations in biophysical properties of the viral membrane is unknown to date. Therefore, we investigated the lifetimes of the probes in naked HIV-1 Gag-based VLPs, or pseudotyped with SARS-CoV-2 WT and delta-Spike protein (nVLP, S-VLP, delta-VLP) (Figure 4A). Similar to the LUV-measurements, the lifetimes were examined in 20 nm windows.

**Figure 4:**
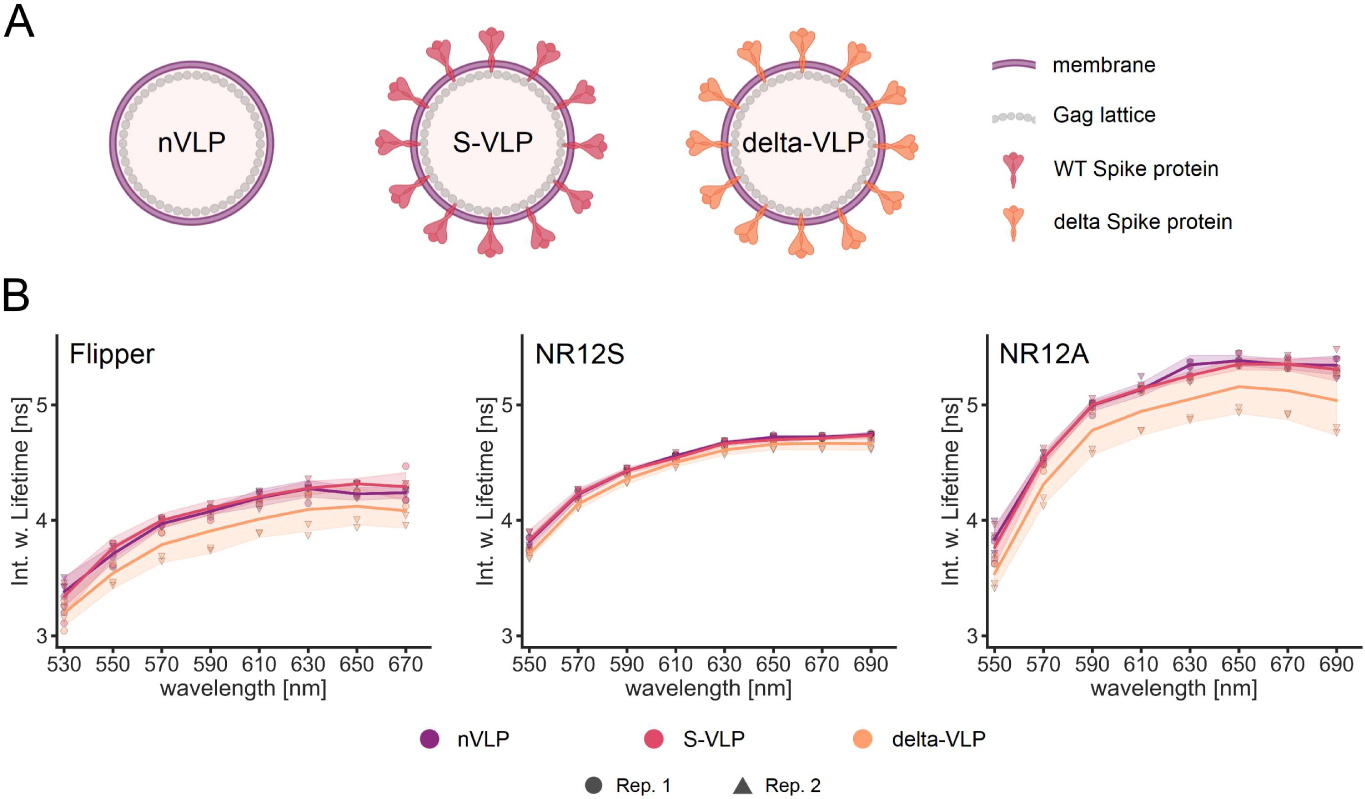
Lifetimes of Flipper, NR12S and NR12A in different VLP species. A| Schematic representation of the SARS-CoV-2 n-VLPs, S-VLPs or delta-VLPs used. B| Spectral fluorescence lifetime measurements of the probes in the VLPs were carried out within 500-700 nm in intervals of 20 nm. Multiexponential curve fitting was performed for the fluorescence decays (for details see Material and Methods). Spectrally resolved intensity weighted lifetime of Flipper (left), NR12S (middle) and NR12A (right) in different VLP species (colour). Line corresponds to the median of individual biological replicates shown with different symbols (n=2). Band corresponds to standard deviation.

Our results indicate that Flipper and NR12A can differentiate membrane properties of delta-VLPs compared to other investigated VLPs by a lifetime shift to lower values. However, both probes exhibit a higher standard deviation of the lifetime in delta-VLPs. The lifetime of NR12S is hardly sensitive to the different VLP types (Figure 4B). The above-reported emission-wavelength dependency is again observed for the VLP measurements of all three probes, while less pronounced compared to the liposomes. Fitting analysis was quite reliable for all probes indicated by χ^2^ values (Supplement Figure 6).

Viral membranes comprise different lipid species and are known to be highly enriched in sphingomyelin and cholesterol (up to 50 %) ^48–51^. In fact, the lifetimes of the probes in the different VLP species are resembling the lifetimes in POPC with 50% cholesterol. Lifetimes of Flipper and NR12A are more sensitive to high cholesterol content than NR12S, indicating that there might be a slightly lower cholesterol content in delta-VLPs, which is not detectable by NR12S. A previous study using NR12S did not observe detectable differences in diffusion or GP value between nVLPs and delta-VLPs ^34^. In addition to cholesterol sensitivity, Flipper also reports on membrane tension, indicating that delta-VLPs might also exhibit distinct membrane tension. Application of membrane tension in single lipid species membranes results in a lower lifetime due to lipid decompression, which gives Flipper more space to relax into its more twisted conformation ^26^. However, viral membranes are heterogenous, in which application of external membrane tension (not caused by increased lipid packing due to cholesterol incorporation) results in higher lifetimes due to membrane reorganization ^26^. Taken together, slight alterations in cholesterol content and/or an increase in membrane tension potentially explain the observed decrease in lifetimes of NR12A and Flipper in delta-VLPs, respectively. Nevertheless, future studies should focus on determination of multiple parameters (e.g. fluidity, tension and charge among others) at once, to better describe the biophysical profiles of viral membranes.

### Which analysis method should be used for NR12S and NR12A: ratiometric GP or fluorescence lifetime?

Given that NR12S and NR12A are mainly used for intensity-based ratiometric analysis (GP), we wanted to compare the resolution power of lifetime vs. GP for all three probes in LUVs of different lipid composition (Figure 5). The wavelengths for GP calculation were chosen to give high GP resolution while maintaining sufficient signal intensity (see Methods). As expected, Flipper shows high sensitivity for different lipid compositions by lifetime analysis, as compared to GP analysis, due to its minor shifts of its emission spectrum (Figure 1A).

**Figure 5:**
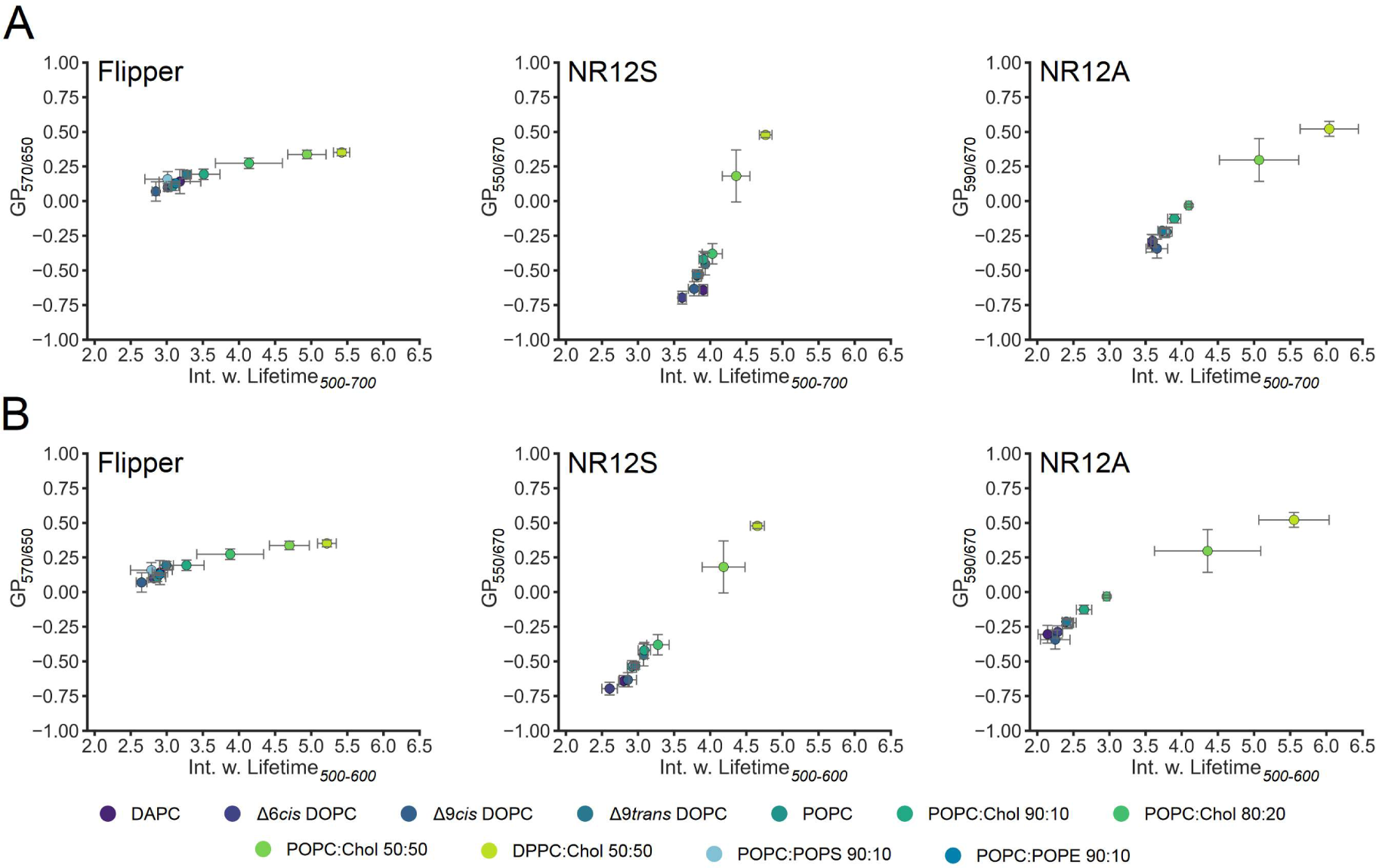
GP analysis vs. lifetime of Flipper, NR12S and NR12A in different lipid environments. Spectral fluorescence lifetime measurements of the probes in LUVs were carried out within 500-700 nm in intervals of 20 nm. Multiexponential curve fitting was performed for the fluorescence decays within A| 500-700 nm or B| 500-600 nm wavelength ranges (for details see Material and Methods). Comparison of resolution power of Flipper (left), NR12S (middle) and NR12A (right) in intensity-based GP analysis vs. lifetime analysis in different lipid environments. Dots correspond to median GP of the respective wavelengths and median mean intensity weighted lifetime. Error bars correspond to the standard deviation. GP and lifetime analysis was performed using the same dataset (for details see Material and Methods).

When the whole spectrum is used for lifetime estimation (mean intensity weighted lifetime within 500-700 nm) for NR12A and NR12S, ratiometric GP-analysis is clearly advantageous in differentiating different lipid compositions over lifetime-based analyses (Figure 5A). However, adjusting the detection window to shorter emission wavelengths (green channel, 500-600 nm), the lifetime resolution especially in more unsaturated, low cholesterol lipid environments almost doubles (Figure 5B). In contrast, the lifetime resolution for Flipper remains unchanged. These results highlight that the suitability of lifetime-based measurements of NR12S and NR12A for membrane fluidity measurements strongly depends on the emission detection settings. Ultimately, using both lifetime and intensity information together, separation of different membrane environments could further improve.

## Conclusion

Our understanding of the influence of biophysical properties on cellular membranes has greatly benefitted from environment-sensitive fluorescent probes. An increasing number of probes have been developed, and are being constantly improved in their functional abilities regarding membrane localization and outer leaflet selectivity ^12,16,17,19^ as well as photostability and brightness allowing even for super-resolution imaging ^18^. Furthermore, their individual sensitivities to varying biophysical properties are being explored ^24,25,27^. Some of these probes have been modified for organelle membrane specificity, allowing the dissection of biophysical properties of different cellular compartments ^28,52^. While many of the solvatochromic probes have been investigated and characterized by spectral intensity-imaging and subsequent GP analysis ^21,25^, the aim of this work was to investigate the probes NR12S and NR12A, alongside Flipper in different lipid environments utilizing FLIM measurements. We explored the sensitivity of the lifetime in low complexity model membrane systems, as well as high complexity live cells and VLPs.

Our work reveals that the lifetimes of NR12S and NR12A are mostly sensitive to larger changes in cholesterol content and saturated vs monounsaturated lipid content, reliably differentiating Lo from Ld phases. Further, the lifetime of NR12A reports on different VLP species. Both probes show a large increase in lifetime with emission wavelength, likely due to solvent relaxation effects and complex photophysical behaviour. The mechanism underlying lifetime shifts in different lipid compositions is most likely also solvent relaxation, as disordered membranes are interspersed with more water molecules facilitating solvent relaxation. Thus resulting in a shorter lifetime for disordered membranes and vice versa for ordered membranes ^37^. The location and orientation of the probes within the membrane also have a big impact on solvent relaxation, as our previous study for NR12S and NR12A applying time-dependent fluorescent shift analyses and atomistic molecular dynamics simulations show ^25^. Flipper is located among the lipid tails within the membrane ^43^, making it less susceptible, but not immune, to solvent relaxation as indicated by its less pronounced emission-wavelength dependency. Using FLIM, we observed that Flipper is primarily sensitive to cholesterol content, can reliably distinguish different membrane phases and reports on different VLP species. The underlying mechanistic principle of variations in the lifetime of Flipper is its planarization upon membrane tension or changes in lipid composition (e.g. cholesterol incorporation) resulting in altered lipid packing ^26,27^. However, in complex heterogenous membranes the impact of either lipid composition or membrane tension is not at all trivial to dissect.

Moreover, we would like to emphasize that although FLIM allows an intensity-independent readout, fluorescence lifetime measurements, subsequent analysis and interpretation with these environment-sensitive membrane probes are not trivial. For reliable fitting of the fluorescence decay a sufficient photon count rate and the adequate laser frequency must be provided to avoid technical pitfalls in experiments. For fluorescent probes with multi-exponential decays, which is true for all three probes used in this study, high photon counts need to be collected for optimal analysis. The technical concerns imply a need for high laser intensities and result in increasing phototoxicity in live samples and bleaching of the probes. Lower intensities generate technical concerns for long acquisition times, which are problematic for moving specimens and probe internalization. To unravel potential small-scale heterogeneities in more complex biological systems, pixel-wise lifetime fitting is required. However, a pixel-wise fit is almost impossible due to the photon budget constraints of these probes, which results in lifetime averaging analysis (image area selection). Fit-independent analysis applying phasor plots can here be explored to group average lifetime components in an image. Furthermore, not all FLIM setups are equipped with pulse pickers allowing to adjust the laser frequency. To allow full exploration of lifetime dynamics across different emission spectra could result in additional need for high-cost equipment, if reliable curve fitting is to be expected. The quantified emission wavelength dependency exhibited by all three membrane probes in this study, indicates possibilities and concerns, especially regarding spectral settings in lifetime measurements. This effect is especially true for NR12S and NR12A as their lifetime resolution is largely increases at shorter wavelengths (500-600 nm) compared to longer wavelengths or the full spectrum. Moreover, our detected wavelength dependency as well as optimum resolution of the lifetime should be investigated and considered for other membrane probes commonly used for lifetime analysis such as Di-4-ANEPPDHQ or Laurdan. Seeing more complex multi-decays of investigated probes at shorter wavelengths, compared to mono-exponential decays at longer wavelengths, must also be used beneficially or with concern.

In summary, we provided an in-depth analysis of the lifetime behaviour of the probes Flipper, NR12S, and NR12A in different lipid environments of model membranes as well as physiological membranes. Further, we emphasized the important factors regarding data acquisition and analysis to be kept in mind when applying these probes in different biological contexts using FLIM.

## Data accessibility

All data will be available upon publication in FigShare.

## Authors’ contribution

F.R.: conceptualization, data curation, formal analysis, investigation, methodology, supervision, validation, visualization, writing—original draft, writing—review and editing;

E.S.: formal analysis, investigation, methodology, writing–original draft

Y.B.U.: formal analysis, investigation, methodology;

J.S: formal analysis, investigation, methodology, supervision, writing–original draft

R.E.: investigation

D.U.: conceptualization, investigation

I.U.: conceptualization, investigation

H.B.: methodology, resources

E.S.: conceptualization, funding acquisition, investigation, methodology, project administration, resources, supervision, validation, writing—original draft, writing—review and editing.

All authors gave final approval for publication and agreed to be held accountable for the work performed therein.

## Acknowledgements

We especially thank the SciLifeLab Advanced Light Microscopy facility and National Microscopy Infrastructure (VR-RFI 2016-00968) for their support on imaging.

---

**Supplement Figure 1: Lifetimes of Flipper, NR12S and NR12A in different lipid environments.** Spectral fluorescence lifetime measurements of the probes in LUVs were carried out within 500-700 nm in intervals of 20 nm. Multiexponential curve fitting was performed for the fluorescence decays (for details see Material and Methods). Spectrally resolved intensity weighted lifetime of Flipper (left), NR12S (middle) and NR12A (right) in different lipid environments investigating cholesterol content (above), double bond position and configuration (middle) and headgroup geometry and charge (below). Line corresponds to the median of individual biological replicates shown with different symbols (n=3 or 4). Band corresponds to standard deviation.

**Supplement Figure 2: Chi-squared values of the multiexponential curve fitting of Flipper, NR12S and NR12A in different lipid environments.** Spectral fluorescence lifetime measurements of the probes in LUVs were carried out within 500-700 nm in intervals of 20 nm. Multiexponential curve fitting was performed for the fluorescence decays (for details see Material and Methods). χ^2^ values serve as indicator for the goodness of the fit and were obtained for each 20 nm interval and are shown for Flipper (left), NR12S (middle) and NR12A (right) in different lipid environments investigating cholesterol content (1^st^ row), saturation index (2^nd^ row) double bond position and configuration (3^rd^ row) and headgroup geometry and charge (4^th^ row). Line corresponds to the median of individual biological replicates shown with different symbols (n=3 or 4).

**Supplement Figure 3: Influence of laser frequency on lifetime analysis**. Spectral fluorescence lifetime measurements of the probes in LUVs were carried out within 500-700 nm in intervals of 20 nm. Multiexponential curve fitting was performed for the fluorescence decays (for details see Material and Methods). The estimated lifetimes of multiexponential curve fitting and the goodness of the fit are evaluated at different laser frequencies: 20, 40 and 80 MHz. A| Spectrally resolved intensity weighted lifetime of NR12A in in Δ9*cis* DOPC (left, blue), POPC (middle, cyan) and DPPC:Chol 50:50 (right, green) at different laser frequencies. The corresponding χ^2^ values serve as indicator for the goodness of the fit and were obtained for each 20 nm interval and are shown below. The line and bar correspond to the median and average, respectively, of two technical replicates. B| Fluorescence decays (grey) and their corresponding fit (green) of NR12S in DPPC:Chol 50:50 at 650 nm at different laser frequencies: 20 MHz (left), 40 MHz (middle) and 80 MHz (right). Corresponding residual counts in artificial units are shown below.

**Supplement Figure 4: Phase selection and Chi-squared values of the multiexponential curve fitting of Flipper, NR12S and NR12A in phase-separated GUVs**. Lifetime measurements in phase-separated GUVs were carried out at 500-600 nm or 600-700 nm emission Multiexponential curve fitting was performed for the fluorescence decays (for details see Material and Methods). A| Overview of manual phase selection procedure in the LAS X software. Liquid disordered (Ld) and liquid-ordered (Lo) phase were selected separately for each phase-separated GUV and each selection was used for lifetime analysis at 500-600 nm and 600-700 nm. B| χ^2^ values serve as indicator for the goodness of the fit and were obtained for Ld and Lo phase and are shown for Flipper (left), NR12S (middle) and NR12A (right) at 500-600 nm and 600-700 nm. Different symbols correspond to individual biological replicates (n=3).

**Supplement Figure 5: Chi-squared values of the multiexponential curve fitting of Flipper, NR12S and NR12A in different cell types.** Lifetime measurements in NRK 52E, U2OS and RBL cells were carried out at 500-600 nm or 600-700 nm emission. Multiexponential curve fitting was performed for the whole-image fluorescence decays (for details see Material and Methods). χ^2^ values serve as indicator for the goodness of the fit and were obtained for different cell types and are shown for Flipper (left), NR12S (middle) and NR12A (right) at 500-600 nm and 600-700 nm. Different symbols correspond to individual biological replicates (n=3).

**Supplement Figure 6: Chi-squared values of the multiexponential curve fitting of Flipper, NR12S and NR12A in different VLP species.** Spectral fluorescence lifetime measurements of the probes in SARS-CoV-2 n-VLPs, S-VLPs or delta-VLPs were carried out within 500-700 nm in intervals of 20 nm. Multiexponential curve fitting was performed for the fluorescence decays (for details see Material and Methods). χ^2^ values serve as indicator for the goodness of the fit and were obtained for each 20 nm interval and are shown for Flipper (left), NR12S (middle) and NR12A (right) in different VLP species. Line corresponds to the median of individual biological replicates shown with different symbols (n=2).

