## Supplementary Information for "Quantifying fluorescence lifetime responsiveness of environment sensitive probes for membrane fluidity measurements"

**Supplement Table 1: Acquisition parameters for all FLIM measurements.**

| | probe | laser intensity | frame repetitions | line repetitions | scan speed | detector | emission |
|---|---|---|---|---|---|---|---|
| LUVs | Flipper (1 ∝M) | 20% | 10 - 20 | 1 | 400 Hz | SMD HyD (10% Gain) photon-counting | 500-700 nm in 20 nm intervals (sequentially) |
| | NR12S (1 ∝M) | 15% | 10 | 1 | | | |
| | NR12A (1 ∝M) | 15% | 10 | 1 | | | |
| | AF488 (1 ∝M) | 2% | 10 | 1 | | | |
| phase-separated GUVs | Flipper (300 nM) | 5% (500-600 nm)<br>10% (600-700 nm) | 1 | 4 | 200 Hz | | 500-600 nm & 600-700 nm (sequentially) |
| | NR12S (100 nM) | 5% | 1 | 2 | | | |
| | NR12A (100 nM) | 5% | 1 | 2 | | | |
| cells | Flipper (1 ∝M) | 3-5% | 1 | 4 | 200 Hz | | 500-600 nm & 600-700 nm (sequentially) |
| | NR12S (1 ∝M) | 1% | 1 | 4 | | | |
| | NR12A (1 ∝M) | 0.2 -1 % | 1 | 4 | | | |
| VLPs | Flipper (300 nM) | 15% | 2 | 16 | 400 Hz | | 500-700 nm in 20 nm intervals (sequentially) |
| | NR12S (100 nM) | 15% | 2 | 16 | | | |
| | NR12A (100 nM) | 15% | 2 | 16 | | | |

**Supplement Table 2: Fitting parameters for all FLIM measurements.**

| | analysis area | laser frequency | fitting range | probe | number of fitting components | environment |
|---|---|---|---|---|---|---|
| **LUVs** | whole image analysis | 20 MHz | 0.2-45 ns for each 20 nm window | Flipper | 2 components: 500-700 nm | all lipid compositions |
| | | | | NR12S | 2 components: 500-620 nm<br>1 component: 620-700 nm | DAPC, Δ6*cis* DOPC, Δ9*cis* DOPC, Δ9*trans* DOPC, POPC, POPC:Chol 90:10, POPC:Chol 80:20, POPS, POPE |
| | | | | | 2 components: 500-600 nm<br>1 component: 600-700 nm | POPC:Chol 50:50, DPPC:Chol 50:50 |
| | | | | NR12A | 2 components: 500-640 nm<br>1 component: 640-700 nm | DAPC, Δ6*cis* DOPC, Δ9*cis* DOPC, Δ9*trans* DOPC, POPC, POPC:Chol 90:10, POPS, POPE |
| | | | | | 2 components: 500-620 nm<br>1 component: 620-700 nm | POPC:Chol 80:20, POPC:Chol 50:50, DPPC:Chol 50:50 |
| | | | | AF 488 | 1 component: 500-700 nm | in water |
| | | | 0.2-45 ns from 500-600 nm or over whole spectrum | Flipper | 2 components | all lipid compositions |
| | | | | NR12S | 2 components | all lipid compositions |
| | | | | NR12A | 2 components | all lipid compositions |
| | | 40 MHz | 0.2-25 ns for each 20 nm window | NR12A | 2 components: 500-640 nm<br>1 component: 640-700 nm | Δ9*cis* DOPC, POPC |
| | | | | | 2 components: 500-620 nm<br>1 component: 620-700 nm | DPPC:Chol 50:50 |
| | | 80 MHz | 0.2-12.5 ns for each 20 nm window | NR12A | 2 components: 500-640 nm<br>1 component: 640-700 nm | Δ9*cis* DOPC, POPC |
| | | | | | 2 components: 500-620 nm<br>1 component: 620-700 nm | DPPC:Chol 50:50 |

| | analysis area | laser frequency | fitting range | probe | number of fitting components | environment |
|---|---|---|---|---|---|---|
| **phase-separated GUVs** | region of interest selection | 20 MHz | 0.2-45 ns for each 100 nm window | Flipper | 2 components: 500-700 nm | Ld and Lo phase |
| | | | | NR12S | 3 components: 500-600 nm<br>1 component: 600-700 nm | Ld phase |
| | | | | | 2 components: 500-600 nm<br>1 component: 600-700 nm | Lo phase |
| | | | | NR12A | 3 components: 500-600 nm<br>2 components: 600-700 nm | Ld phase |
| | | | | | 3 components: 500-600 nm<br>1 component: 600-700 nm | Lo phase |
| **cells** | whole image analysis | 20 MHz | 0.2-45 ns for each 100 nm window | Flipper | 3 components: 500-700 nm | all cell types |
| | | | | NR12S | 3 components: 500-600 nm<br>2 components: 600-700 nm | all cell types |
| | | | | NR12A | 3 components: 500-600 nm<br>1 component: 600-700 nm | all cell types |
| **VLPs** | whole image analysis | 20 MHz | 0.2-45 ns for each 20 nm window | Flipper | 2 components: 500-700 nm | all VLPs |
| | | | | NR12S | 2 components: 500-620 nm<br>1 component: 620-700 nm | all VLPs |
| | | | | NR12A | 2 components: 500-620 nm<br>1 component: 620-700 nm | all VLPs |

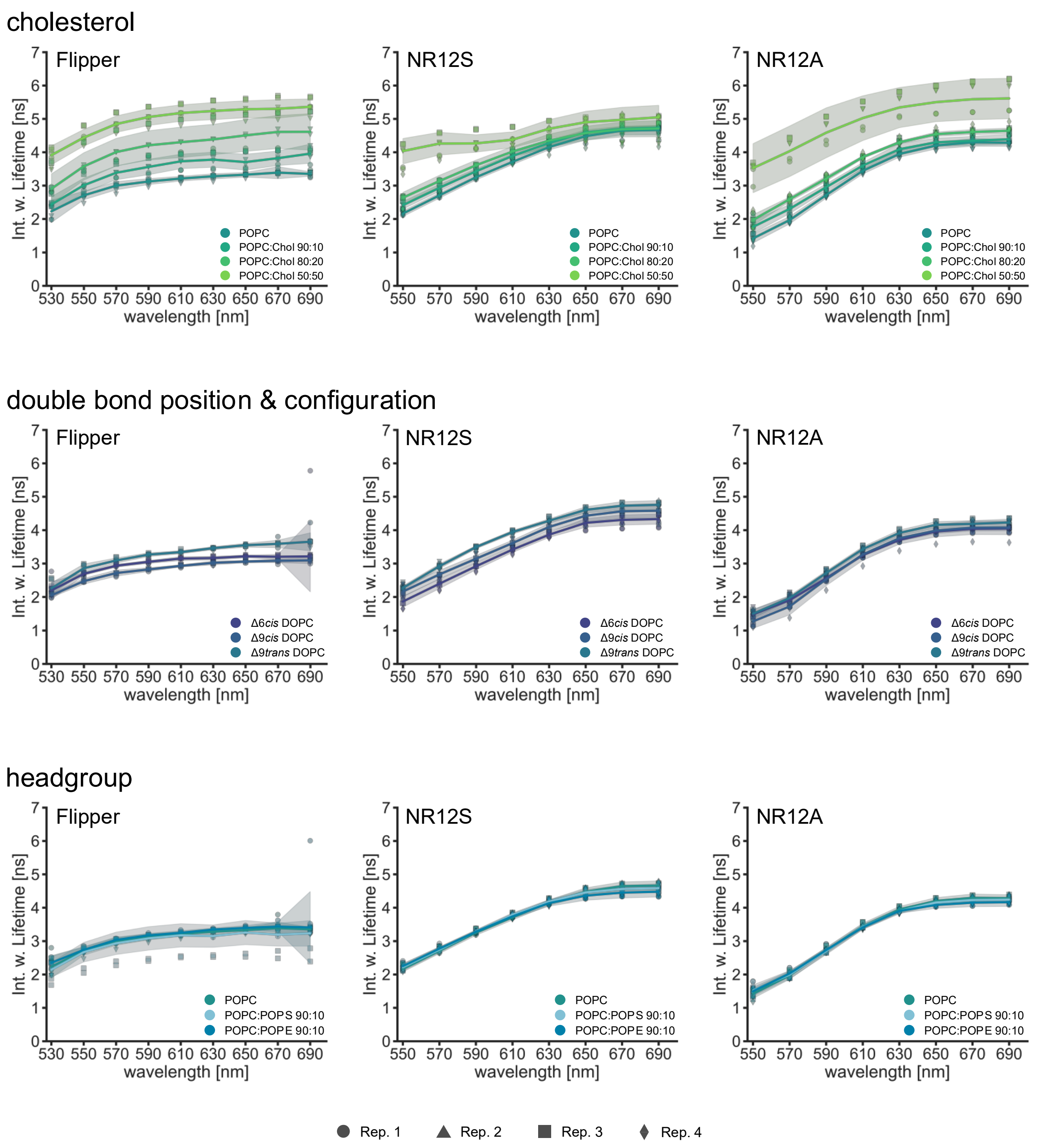


**Supplement Figure 1: Lifetimes of Flipper, NR12S and NR12A in different lipid environments.** Spectral fluorescence lifetime measurements of the probes in LUVs were carried out within 500-700 nm in intervals of 20 nm. Multiexponential curve fitting was performed for the fluorescence decays (for details see Material and Methods). Spectrally resolved intensity weighted lifetime of Flipper (left), NR12S (middle) and NR12A (right) in different lipid environments investigating cholesterol content (above), double bond position and configuration (middle) and headgroup geometry and charge (below). Line corresponds to the median of individual biological replicates shown with different symbols (n=3 or 4). Band corresponds to standard deviation.

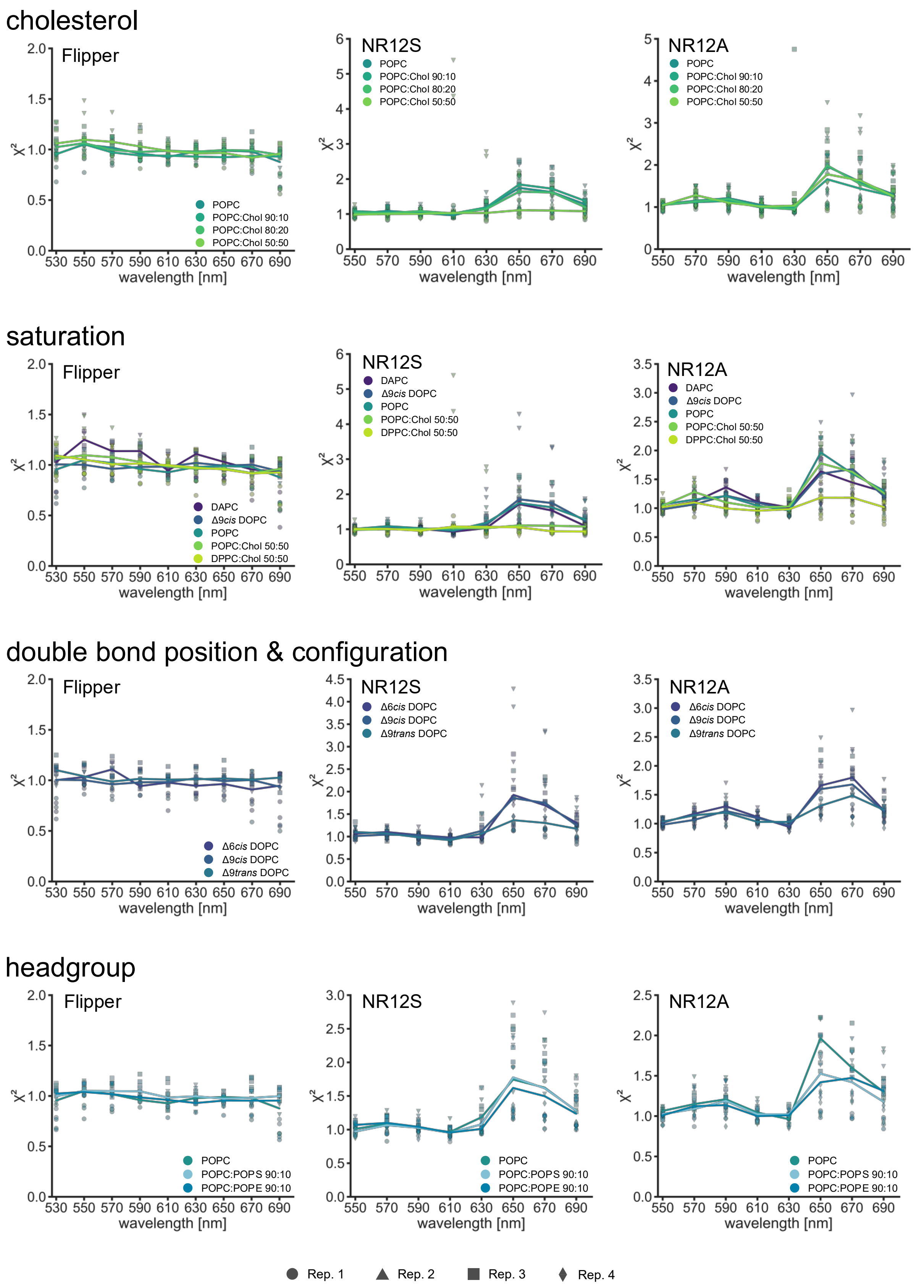


**Supplement Figure 2: Chi-squared values of the multiexponential curve fitting of Flipper, NR12S and NR12A in different lipid environments.** Spectral fluorescence lifetime measurements of the probes in LUVs were carried out within 500-700 nm in intervals of 20 nm. Multiexponential curve fitting was performed for the fluorescence decays (for details see Material and Methods). $\chi^2$ values serve as indicator for the goodness of the fit and were obtained for each 20 nm interval and are shown for Flipper (left), NR12S (middle) and NR12A (right) in different lipid environments investigating cholesterol content (1$^{st}$ row), saturation index (2$^{nd}$ row) double bond position and configuration (3$^{rd}$ row) and headgroup geometry and charge (4$^{th}$ row). Line corresponds to the median of individual biological replicates shown with different symbols (n=3 or 4).

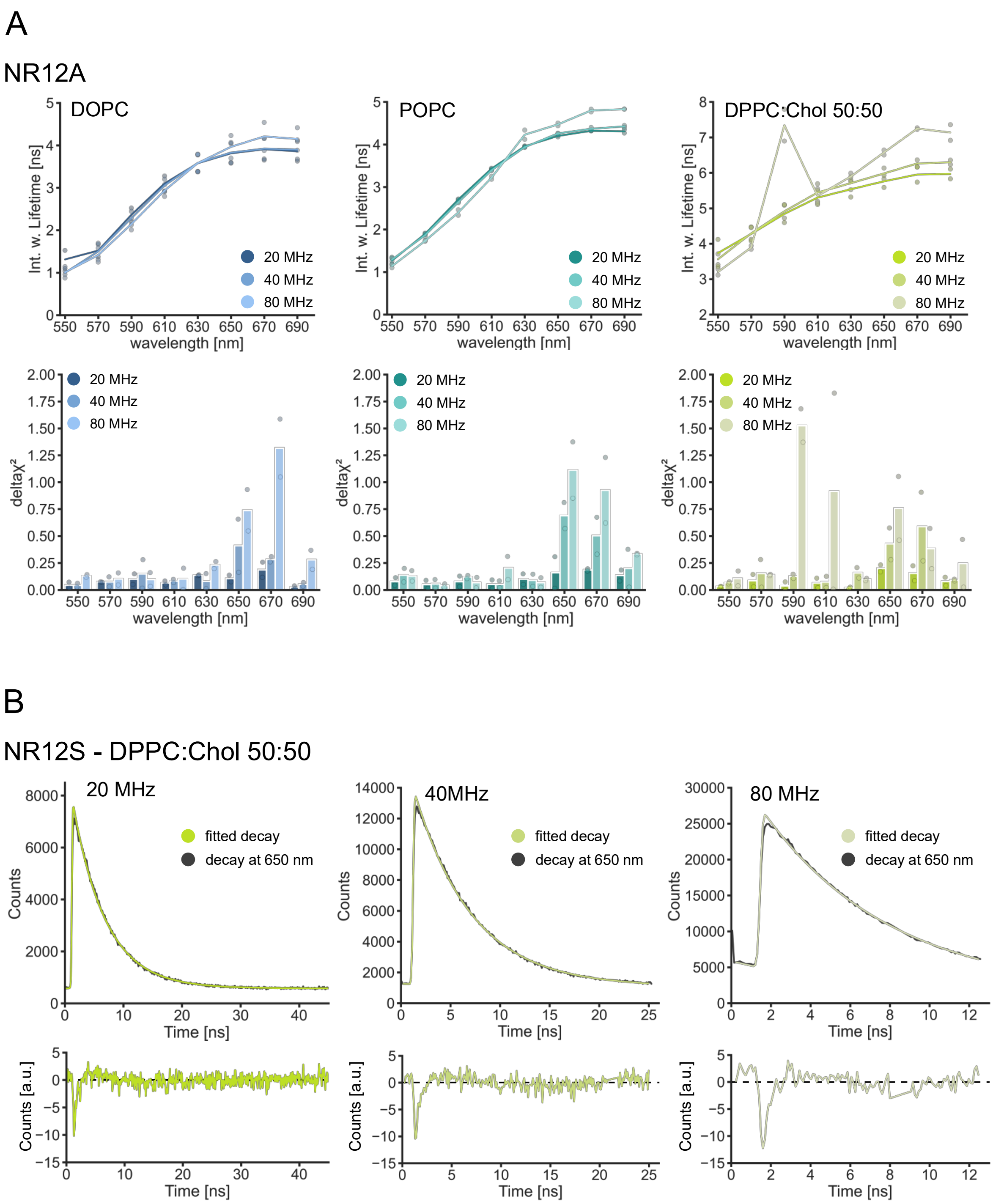


**Supplement Figure 3: Influence of laser frequency on lifetime analysis**. Spectral fluorescence lifetime measurements of the probes in LUVs were carried out within 500-700 nm in intervals of 20 nm. Multiexponential curve fitting was performed for the fluorescence decays (for details see Material and Methods). The estimated lifetimes of multiexponential curve fitting and the goodness of the fit are evaluated at different laser frequencies: 20, 40 and 80 MHz. A| Spectrally resolved intensity weighted lifetime of NR12A in in Δ9*cis* DOPC (left, blue), POPC (middle, cyan) and DPPC:Chol 50:50 (right, green) at different laser frequencies. The corresponding $\chi^2$ values serve as indicator for the goodness of the fit and were obtained for each 20 nm interval and are shown below. The line and bar correspond to the median and average, respectively, of two technical replicates. B| Fluorescence decays (grey) and their corresponding fit (green) of NR12S in DPPC:Chol 50:50 at 650 nm at different laser frequencies: 20 MHz (left), 40 MHz (middle) and 80 MHz (right). Corresponding residual counts in artificial units are shown below.

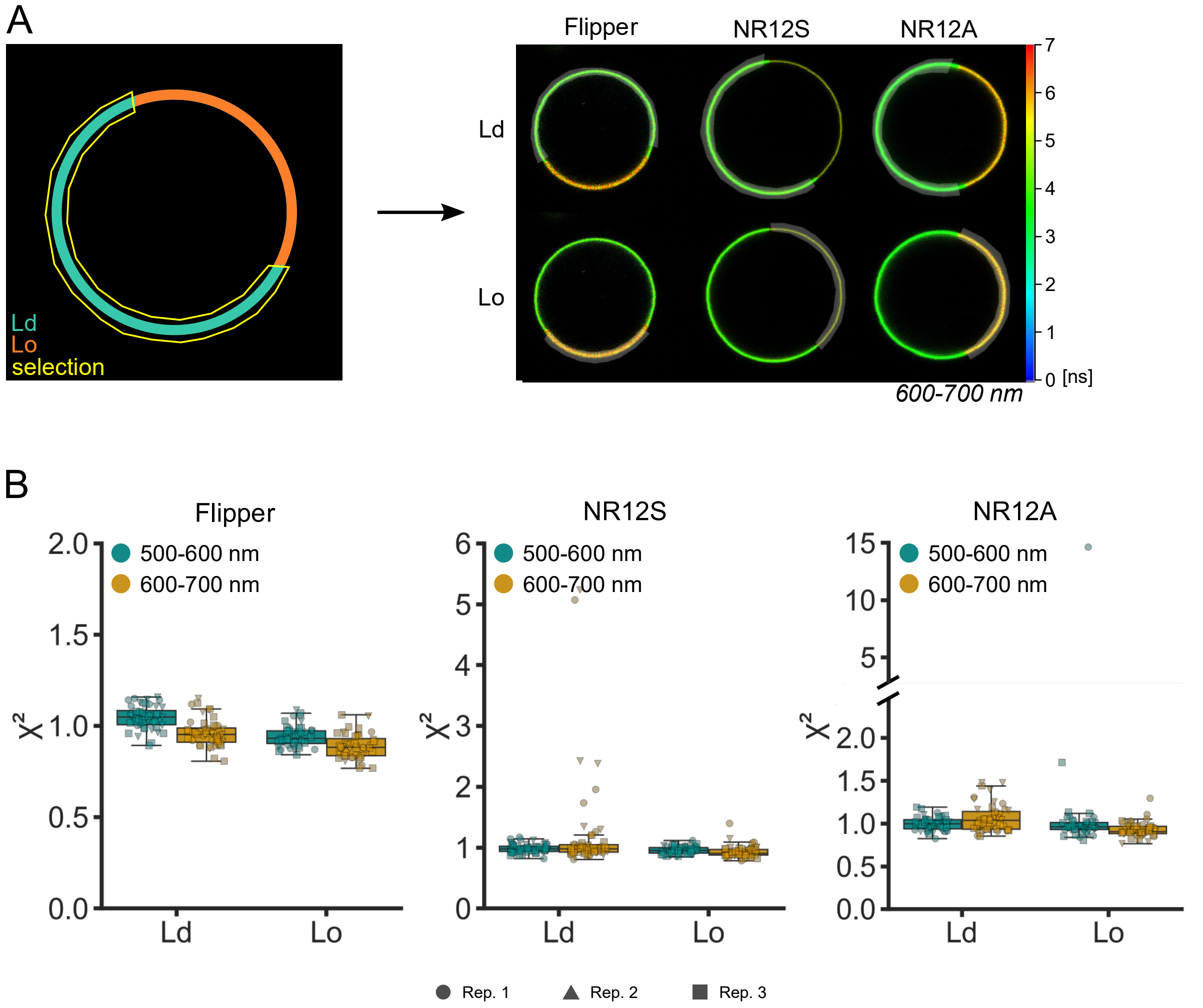


**Supplement Figure 4: Phase selection and Chi-squared values of the multiexponential curve fitting of Flipper, NR12S and NR12A in phase-separated GUVs**. Lifetime measurements in phase-separated GUVs were carried out at 500-600 nm or 600-700 nm emission. Multiexponential curve fitting was performed for the fluorescence decays (for details see Material and Methods). A| Overview of manual phase selection procedure in the LAS X software. Liquid disordered (Ld) and liquid-ordered (Lo) phase were selected separately for each phase-separated GUV and each selection was used for lifetime analysis at 500-600 nm and 600-700 nm. B| $\chi^2$ values serve as indicator for the goodness of the fit and were obtained for Ld and Lo phase and are shown for Flipper (left), NR12S (middle) and NR12A (right) at 500-600 nm and 600-700 nm. Different symbols correspond to individual biological replicates (n=3).

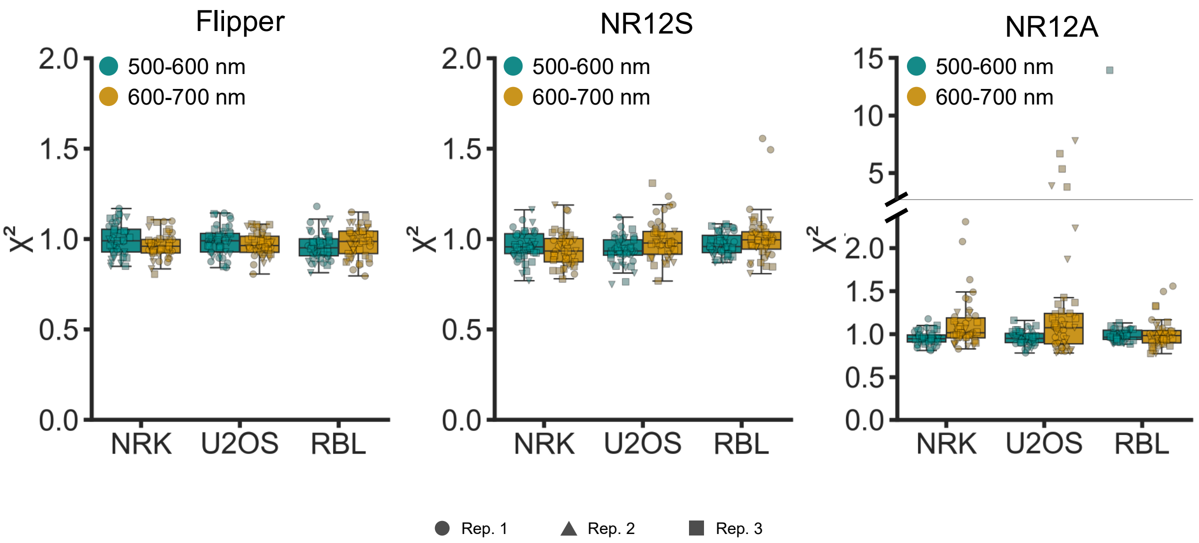


**Supplement Figure 5: Chi-squared values of the multiexponential curve fitting of Flipper, NR12S and NR12A in different cell types.** Lifetime measurements in NRK 52E, U2OS and RBL cells were carried out at 500-600 nm or 600-700 nm emission. Multiexponential curve fitting was performed for the whole-image fluorescence decays (for details see Material and Methods). $\chi^2$ values serve as indicator for the goodness of the fit and were obtained for different cell types and are shown for Flipper (left), NR12S (middle) and NR12A (right) at 500-600 nm and 600-700 nm. Different symbols correspond to individual biological replicates (n=3).

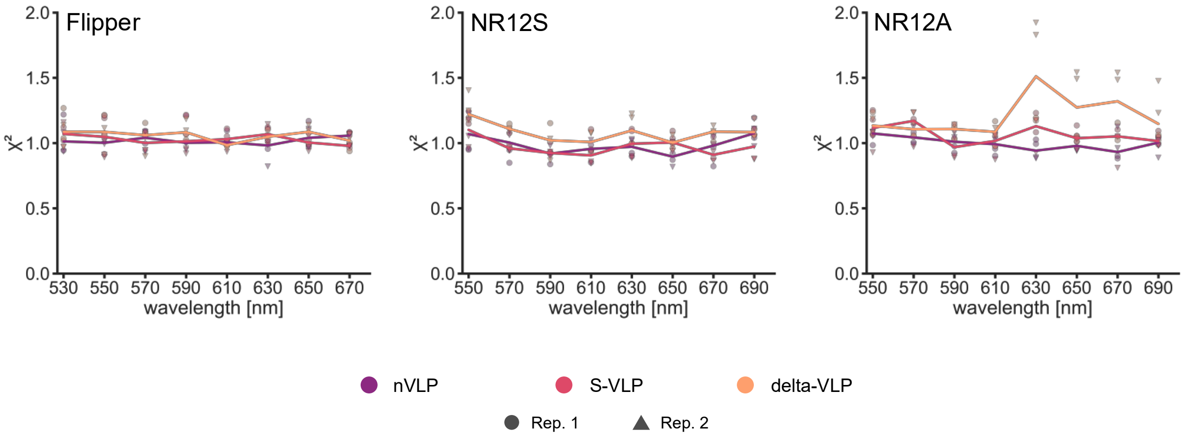


**Supplement Figure 6: Chi-squared values of the multiexponential curve fitting of Flipper, NR12S and NR12A in different VLP species.** Spectral fluorescence lifetime measurements of the probes in SARS-CoV-2 n-VLPs, S-VLPs or delta-VLPs were carried out within 500-700 nm in intervals of 20 nm. Multiexponential curve fitting was performed for the fluorescence decays (for details see Material and Methods). $\chi^2$ values serve as indicator for the goodness of the fit and were obtained for each 20 nm interval and are shown for Flipper (left), NR12S (middle) and NR12A (right) in different VLP species. Line corresponds to the median of individual biological replicates shown with different symbols (n=2).
